# Vascular Endothelial Growth Factor as an Immediate-Early Activator of UV-induced Skin Injury

**DOI:** 10.1101/2020.07.13.198549

**Authors:** Stella P. Hartono, Victoria M. Bedell, Sk. Kayum Alam, Madelyn O’Gorman, MaKayla Serres, Stephanie R. Hall, Krishnendu Pal, Rachel A. Kudgus, Priyabrata Mukherjee, Davis M. Seelig, Alexander Meves, Debabrata Mukhopadhyay, Stephen C. Ekker, Luke H. Hoeppner

## Abstract

The negative health consequences of acute ultraviolet (UV) exposure are evident, with reports of 30,000 emergency room visits annually to treat the effects of sunburn in the United States alone. Acute effects of sunburn include erythema, edema, and severe pain, and chronic overexposure to UV radiation can lead to skin cancer. While the pain associated with the acute effects of sunburn may be relieved by current interventions, existing post-sunburn treatments are not capable of reversing the cumulative and long-term pathological effects of UV exposure, an unmet clinical need. Here we show that activation of the vascular endothelial growth factor (VEGF) pathway is a direct and immediate consequence of acute UV exposure, and activation of VEGF signaling is necessary for the initiating the acute pathological effects of sunburn. In UV-exposed human subjects, VEGF signaling is activated within hours. Topical delivery of VEGF pathway inhibitors, targeted against the ligand VEGF-A (gold nanoparticles conjugated with anti-VEGF antibodies) and small molecule antagonists of VEGF receptor signaling, prevent the development of erythema and edema in UV-exposed mice. Collectively, these findings suggest targeting VEGF signaling may reduce the subsequent inflammation and pathology associated with UV-induced skin damage, which reveals a new post-exposure therapeutic window to potentially inhibit the known detrimental effects of UV on human skin.

---

Skin is our largest organ and is critical for exposure to the environment and for social interaction. Ultraviolet (UV) radiation from the sun elicits dramatic acute and chronic effects. Approximately 30,000 Americans are admitted to the emergency room annually to treat the effects of sunburn, including erythema, edema and severe pain (1). Chronic UV exposure induces decreased skin elasticity, promotes increased wrinkling (2, 3), and is associated with the development of skin cancers (4, 5). A history of severe sunburns has been associated with increased risks for all types of skin cancer, including melanoma, squamous cell carcinoma, and basal cell carcinoma (6). While the increasing incidence of these three types of skin cancer has resulted in a greater focus on sun exposure habits (7), one-third of Americans report experiencing sunburn within the past year (8). Current sunburn treatments, such as non-steroidal anti-inflammatory medications, aloe vera gels, and systemic and topical corticosteroids, may provide immediate pain relief but do not reverse the cumulative and chronic pathological effects of UV exposure (9). Thus, viable post-exposure sunburn treatments represent an unmet clinical need.

In addition to DNA damage (10), UV exposure causes immunomodulation through multiple molecular mechanisms in both the innate and adaptive immune systems. Acute inflammation is a well-established hallmark of the human body’s response to sunburn (11). However, the immediate pathological effects of sunburn (i.e. pain, itching, swelling, redness, skin that feels hot to the touch, etc.) can occur hours before the acute inflammatory response. Vascular endothelial growth factor (VEGF) (12) / vascular permeability factor (VPF) (13) has been implicated in the initial skin response to UV exposure (14, 15). VEGF is upregulated following UV exposure in mice, and increased VEGF causes sensitization to UV and increased severity of sunburn, including greater vascularity and edema (14, 15). However, the importance of VEGF activation in response to acute UV-induced skin damage remains underappreciated.

Here we tested the hypothesis that activation of the VEGF pathway is a direct and immediate consequence of acute UV exposure and is essential for the initiation of the pathological effects of sun exposure in the epidermis. We propose that VEGF induction causes edema and the related immediate impact on the skin, while working concomitantly with reactive oxygen species (ROS) to induce inflammation. Specifically, we show that VEGF signaling is activated within hours of solar simulated light exposure in human subjects. Using a mouse post-UV exposure model, we show that topical delivery of VEGF pathway inhibitors, targeted against the ligand VEGF-A (gold nanoparticles conjugated with anti-VEGF antibodies; aVGNPs) and small molecule antagonists of VEGF receptor signaling, can dramatically prevent the subsequent development of erythema and edema associated with acute sunburn. Taken together, these results demonstrate a novel and clinically relevant method of skin injury treatment following UV exposure, highlighting the understudied role of the VEGF/VPF pathway in both acute and chronic sun damage.

We assessed whether activation of the VEGF pathway is a direct and immediate consequence of acute UV exposure in human subjects exposed to solar simulated UV light. We performed immunostaining for total VEGFR-2 protein and phosphorylation of VEGFR-2 at Y1175 (pVEGFR-2) using skin tissue specimens obtained from four human subjects exposed to a 2.5 minimal erythema dose (MED) of solar-simulated UV light (**Figure 1**). We observed no detectable pVEGFR-2 immunostaining at 0, 5, and 60 minutes following UV exposure. At 5 hours post-UV exposure, mild to moderate pVEGFR-2 was observed within the stratum basale of the epidermis. By 24 hours post-UV exposure, similar pVEGFR-2 was also detectable, but at a lesser frequency (**Figure 1, Supplementary Table 1**). Strong total VEGFR-2 staining was present within the dermis of human skin at all time points, while greater epidermal total VEGFR-2 expression positively correlated with increased duration of solar-simulated UV exposure (**Figure 1, Supplementary Table 1**). Taken together, these results suggest VEGF signaling is activated within 5 hours of UV exposure in humans.

**Figure 1:**
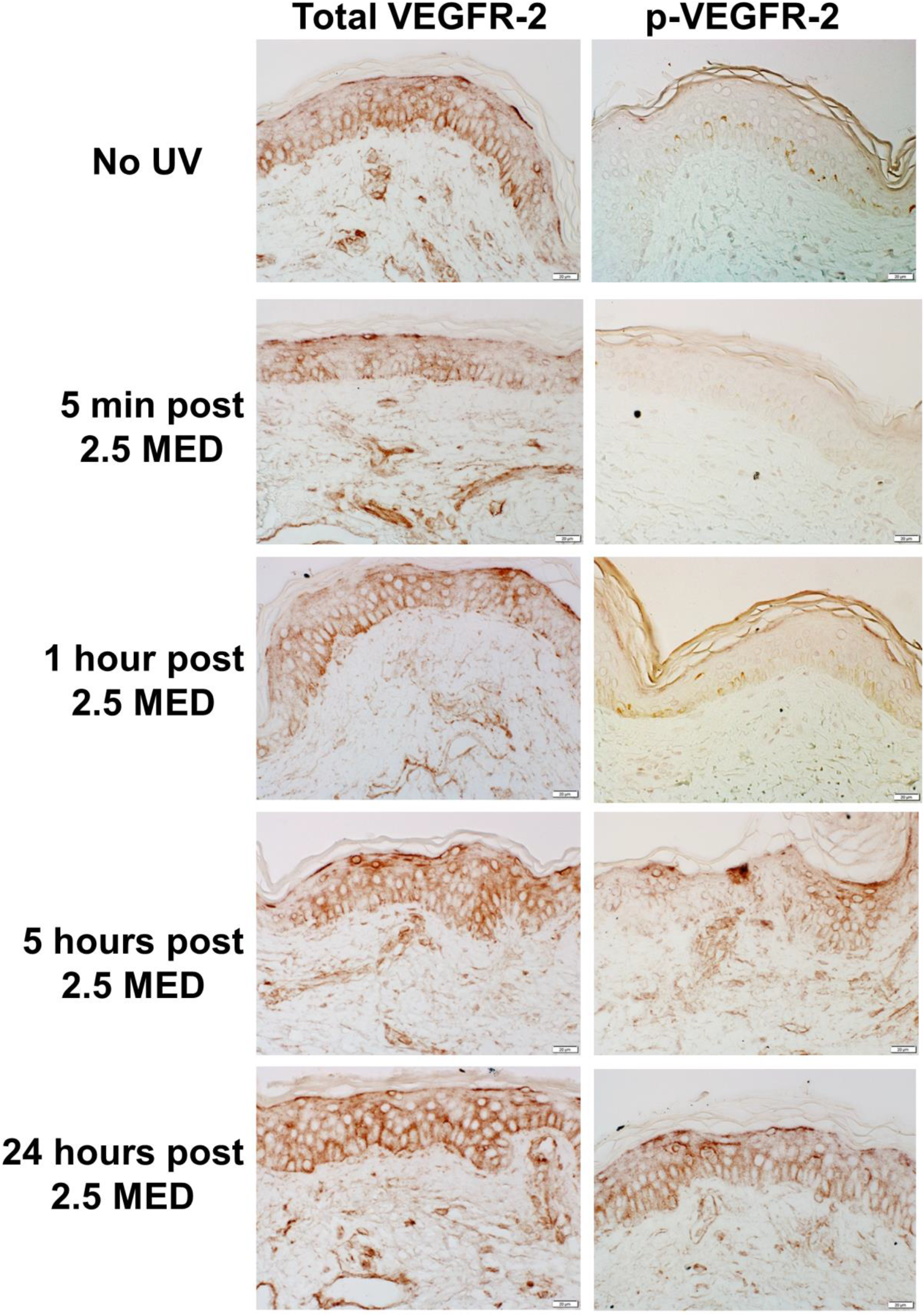
UV exposure activates VEGF signaling in human subjects. Skin tissue specimens collected from human subjects (n=4) at 5 minutes, 1 hour, 5 hours, and 24 hours post-exposure to 2.5 MED solar-simulated UV light were immunostained using antibodies that detect total VEGFR-2 protein (left column) and VEGFR-2 phosphorylated at its Y1175 residue (p-VEGFR-2; right column). Representative images are shown. Scale bar indicates 20 μm.

Exposure to ultraviolet B (UVB) irradiation induces skin alterations such as erythema, dilation of dermal blood vessels, vascular hyper-permeability, and epidermal hyperplasia, which comprises acute photo damage. To determine the minimal erythema dose (MED) required to induce these pathological features of acute skin damage, various doses of UVB were applied to flank skin of immunocompetent, hairless SKH1 mice. Erythema was observed in mice by 48 hours, following exposure to 0.144 J/cm^2^ or greater UVB. We performed qPCR to determine level of VEGF expression in mRNA derived from mice 48 hours after exposure to various doses of UVB. We observed dose dependent increases in the skin starting at 0.144 J/cm^2^ UVB **(Figure 2A)**.

**Figure 2:**
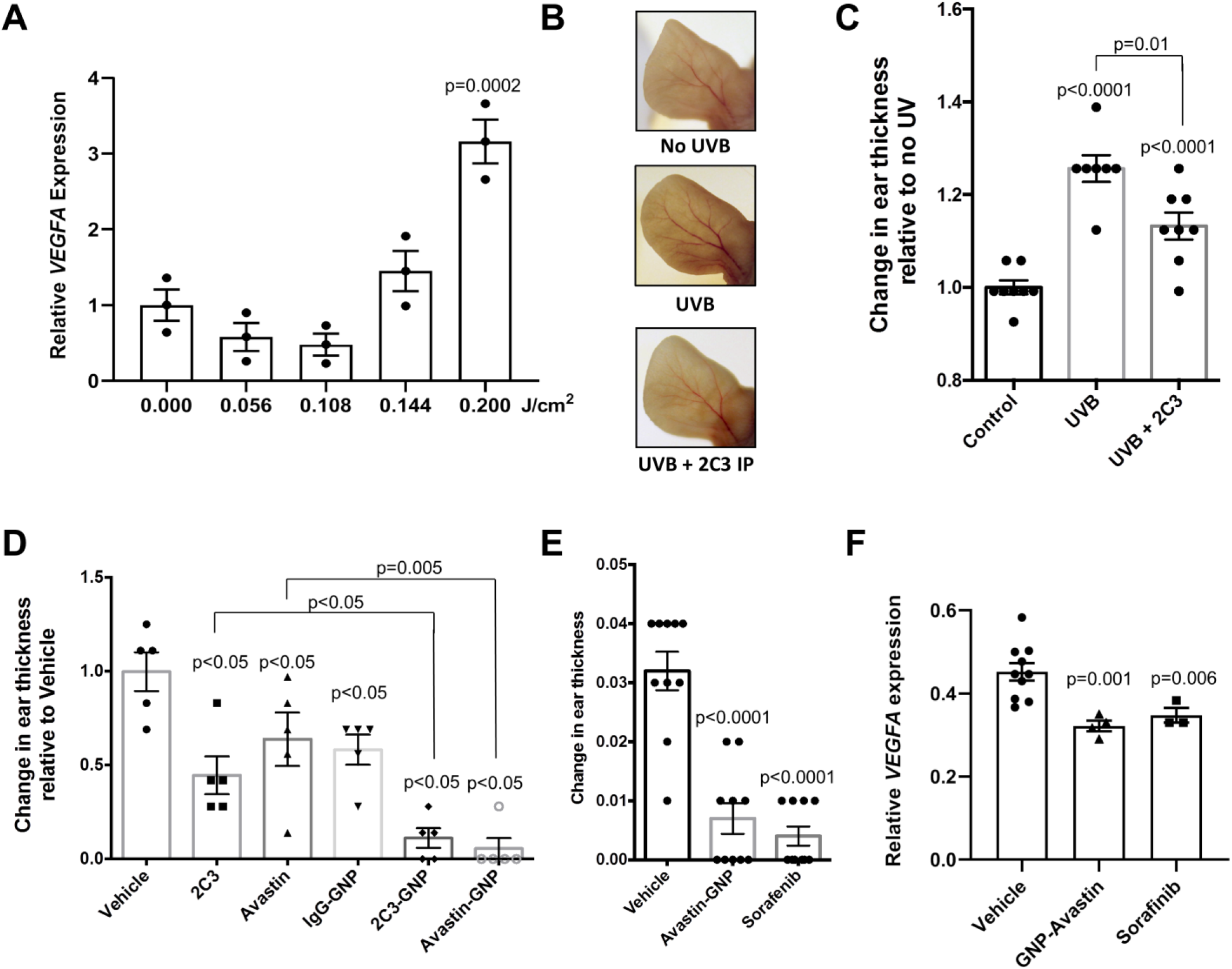
Topical inhibition of VEGF signaling reduces UV-induced skin damage in mice. (**a**) Quantitative PCR was used to measure induction of VEGF transcript in mouse skin at 48 hours post-exposure to various indicated doses of UVB (n=3 mice). (**b, c**) Ears of mice (2 separate experiments, n=4 mice per group) were exposed to a single 1.0 MED UVB exposure of 0.144 J/cm^2^ after treatment with 50 μg VEGF-neutralizing antibody 2C3 or negative control PBS by intraperitoneal injection 24 hours prior. Representative images are shown (**b**). Extent of ear edema was determined by measuring ear thickness 48 hours post radiation. (**c**). (**d**) Ears of mice (n=5 mice per group) were exposed once to 1.0 MED of UVB radiation at 0.144 J/cm^2^. Indicated topical treatments (25 μg per mouse; 12.5 μg per ear) or control vehicle (Vanicream) were applied two hours post-radiation. Extent of ear edema was determined by measuring the change in ear thickness 48 hours post-radiation. **(e, f)** Topical administration of GNP-VEGF antibody (25 ug per mouse; 12.5 μg per ear) or small molecular inhibitor of VEGF signaling, sorafenib (25 ug per mouse; 12.5 μg per ear) reduces edema (2 separate experiments, n=5 mice per group). All treatments were applied 2 hours following 1.0 MED UVB exposure of 0.144 J/cm^2^ UVB. Ear thickness was measured 2 days after UVB exposure. Quantitative PCR was used to measure induction of VEGF transcript in mouse ear skin following UV exposure and treatment in a subset of mice (**f**).

To determine whether VEGF induction is necessary for the development of acute photo damage, we treated mice with an intraperitoneal injection of 50 μg VEGF antibody 2C3 and exposed their ears to 1.0 MED. Along with erythema and increased vascularity, UVB-exposed, untreated mice showed signs of edema as evidenced by increased ear thickness. Intraperitoneal administration of 2C3 decreases blood vessel dilation **(Figure 2B)**, reduces edema as shown by 50% decrease in ear thickness **(Figure 2C),** and reduces angiogenesis as shown by decreased CD31-positive endothelial cells in UVB-exposed mouse ears (**Supplemental Figure 1**). These findings suggest that VEGF is induced early and necessary for the development of acute photo damage.

We investigated the efficacy of topical administration of GNP-conjugated VEGF antibody in preventing acute photo damage after UVB exposure. As we intended this as a treatment instead of prophylactic measures, we applied the topical treatment 2 hours after exposing healthy mice ears to UVB. We showed that topical administration of GNP conjugated to two different VEGF antibodies (2C3 or Avastin) is superior in reducing edema compared to topical VEGF antibody or GNP alone **(Figure 2D)**. Topical administration of sorafenib, a small molecule VEGFR-2 tyrosine kinase inhibitor, similarly reduced mouse ear thickness caused by UVB exposure **(Figure 2E)**. Quantitative PCR of skin tissue RNA confirmed that topical administration of sorafenib or GNP conjugated VEGF antibody reduced VEGF transcript levels in UVB exposed mice **(Figure 2F)**. These results suggest topical inhibition of VEGF signaling after UVB exposure is effective in preventing development of acute photo damage. To our knowledge, this is the first study demonstrating that acute UVB-induced skin injury can be prevented following exposure.

## DISCUSSION

Solar UV radiation is subdivided into three categories, UVA, UVB, and UVC (16). Most UVB rays penetrate the epidermis or upper region of the dermis (17). UVB light is 1,000-10,000 times more carcinogenic than UVA radiation measured by DNA damage and erythema (18). Our studies focus on the molecular initiation of UVB-induced skin damage and its associated pathology, and suggest that topical inhibition of VEGF/VPF signaling reduces photo damage in vivo which occurs well before the activation of the innate immune response.

We propose that early activation of VEGF signaling is a direct and immediate response to acute UVB exposure that leads to edema and related pathological effects on the skin. We demonstrate through immunostaining that VEGFR-2 activation via phosphorylation of the Y1175 residue in the epidermis of human subjects exposed to solar-simulated UV rays peaks at 5 hours post-exposure. Reverse phase protein arrays using similar human specimens suggested that VEGFR-2 Y996 was increased at 1 hour and 5 hours post-UV exposure (19). While the functional consequences of VEGFR-2 Y996 are unknown, this residue serves as a docking site for SH2, SH3, or PTB domain-containing proteins to convey downstream signaling (20, 21). VEGFR-2 Y1175 phosphorylation stimulates the PLCγ-ERK1/2 cascade, regulates Ca2+ signaling as well as cellular survival and proliferation (22). Human keratinocytes and epidermis express VEGF receptors and co-receptors, and autocrine VEGF/VEGFR-2 signaling is activated in response to moderate UVB irradiation (23, 24). VEGF is produced through distinct mechanisms by UVB and ROS (25), as UVB has been shown to activate VEGF in the presence of antioxidants (26). UVB and inflammatory mediator PGE2 directly upregulate VEGF in human dermal fibroblasts and indirectly elevate VEGF in human epidermal keratinocytes (27). Potentiation of the VEGF/VEGFR-2-mediated angiogenic response induced by UVB in VEGF transgenic mice stimulated UV-induced cutaneous skin damage, yet did not contribute to wound healing and repair mechanisms (14).

We show that VEGF-A is induced immediately following acute UVB exposure in mice. The acute UVB-induced edema, erythema, and increased vascularity can be substantially reduced by disrupting VEGF signaling via systemic or topical administration of anti-VEGF antibody. GNPs possess intrinsic anti-angiogenic properties due to selective inhibitory interactions with heparin-binding growth factors, such as VEGF and bFGF (28). Indeed, topical application of GNP conjugated to VEGF antibody synergistically reduces acute UVB-induced edema in mice. This is supported by studies demonstrating that UVB irradiation induces an angiogenic switch mediated by upregulation of VEGF and that elevation of VEGF increases photosensitivity (14, 15, 29). This effect is mediated through activation of VEGFR-2 as topical administration of sorafenib produces similar improvement. VEGF/VEGFR-2 transduces signals through JAK/STAT proteins (30), and inhibition of JAK2/STAT3-dependent autophagy with sanshool exhibited a photoprotective effect in human dermal fibroblasts and hairless mice exposed to UVB irradiation (31). Our findings suggest VEGF/VPF signaling is an immediate-early activator of UV-induced skin injury and plays an integral role in the associated pathology, which occurs well before the innate immune response. Thus, targeting VEGF/VEGFR-2 signaling reduce the subsequent inflammation and pathology associated with UV-induced skin damage.

## MATERIALS AND METHODS

### Mice

Six- to eight-week-old female SKH1-Elite hairless mice were purchased (Stain Code: 477, Charles River). All mice were housed in a temperature-controlled room with alternating 12 h light/dark cycles, allowed one week to acclimate to their surroundings, and fed a standard diet. All animal work was conducted in accordance with protocols approved by Institutional Animal Care and Use Committee at Mayo Clinic and University of Minnesota.

### Gold-nanoparticle antibody conjugation

Naked gold nanoparticles (GNPs) were synthesized by adding 500 ml of an aqueous solution containing 43 mg of sodium borohydride (Sigma Aldrich) to 1000 ml of 0.1 mM HAuCl4 (Sigma Aldrich) solution under constant stirring, overnight at room temperature. The desired antibody (800 μg of 2C3, Avastin, or IgG) was suspended in 1 ml of molecular biology grade water and added dropwise to 200 ml of naked GNP solution under constant stirring at ambient temperature for two hours. The mixture was centrifuged at 22,000 rpm in a Beckman Coulter ultracentrifuge at 4°C for 65 minutes twice to separate the GNP-antibody conjugate from naked antibody. The supernatant was removed and the conjugated GNP-antibody pellet was suspended in molecular biology grade water to the desired concentration.

### Preparation of gold-nanoparticle antibody cream

Vanicream (Pharmaceutical Specialties, Inc.) was mixed with an equal volume of GNP-antibody conjugate. The typical dose of GNP-antibody administered topically to each mouse was 25 μg unless indicated otherwise. VEGF neutralizing antibodies used were 2C3 (Peregrine Pharmaceuticals) and bevacizumab (Avastin®, Genentech).

### UV Irradiation Studies

Six- to eight-week-old female SKH1-Elite hairless mice were exposed to graded doses of a single UVB irradiation using fluorescent lamps. The height of the lamps was adjusted to deliver 0.82 mW/cm^2^ at the dorsal skin surface. The minimal erythema dose (MED) was determined by irradiation of eight 1 cm^2^ areas on the skin on the front of mice with seven graded doses of UVB irradiation ranging from 0.056 J/cm^2^ to 0.4 J/cm^2^ as well as a sham irradiation. Erythema formation was evaluated after 48 hours by two independent observers. To determine systemic effect of VEGF inhibition on acute UVB skin injury, mice were treated with 50 μg VEGF-neutralizing antibody 2C3 by intraperitoneal injection 24 hours prior to exposure of 0.144 J/cm^2^ of UVB radiation to the ears. Extent of ear edema was determined by measuring ear thickness as previously described (14). Samples of dorsal skin and ears were snap-frozen in liquid nitrogen or fixed in formaldehyde.

To evaluate effectiveness of topical treatment on acute UVB exposure, ears of Six- to eight-week-old female C57BL/6 (n=5 per group) were exposed to one dose of 0.144 J/cm^2^ UVB radiation. Topical treatments (25 ug/mice mixed in 1 mL of Vanicream) or control vehicle (Vanicream) were applied two hours post-radiation. Extent of ear edema was determined by measuring the change in ear thickness 48 hours post-radiation. Samples of ears were snap-frozen in liquid nitrogen or fixed in formaldehyde.

### Immunohistochemistry staining

Human skin tissue specimens were collected from each of four human subjects at 5 minutes, 1 hour, 5 hours, and 24 hours post-exposure to 2.5 MED solar-simulated UV light at the University of Arizona in accordance with Institutional Review Board approval and informed written consent of all study participants (19). Formalin-fixed, paraffin embedded, serially sectioned specimens mounted on glass slides were a generous gift from Dr. Clara Curiel-Lewandrowski at the University of Arizona. Slides were immunostained using antibodies that recognize total VEGFR-2 (#2479, Cell Signaling Technology) and phosphorylation of VEGFR-2 at Y1175 (#2478, Cell Signaling Technology) as previously described (32). Pathological review and analysis of the immunostaining were performed by a board-certified pathologist (D.M.S.) at the University of Minnesota, and histological findings have been reported in Figure 1 and Supplementary Table 1.

Tissue was harvested from the ears of mice exposed to a single 1.0 MED UVB exposure of 0.144 J/cm^2^, preceded by an intraperitoneal injection with 50 μg VEGF-neutralizing antibody 2C3 or negative control PBS 24 hours before UVB exposure. Formalin-fixed, paraffin embedded, serially sectioned specimens mounted on glass slides were immunostained using a polyclonal CD31 antibody (sc-1506, Santa Cruz Biotechnology). Following analysis of the immunostaining, representative histological images have been reported in Supplementary Figure 1.

### Quantitative PCR Analysis

Purified RNA was isolated from mouse skin using the RNeasy Plus Mini Kit (Qiagen) after pulverization of the frozen skin tissue and homogenization with the QIAshredder system (Qiagen). Quantitative PCR was performed using the QuantiTech SYBR Green RT-PCR kit (Qiagen) per the manufacturer’s instructions. Briefly, RNA (50 ng) was added to 30 μl reactions with QuantiTech SYBR Green RT master mix, QuantiTech RT mix, and 0.5 pmol/μl of each of the oligonucleotide primers.

Total RNA was isolated from frozen mouse skin tissues using the RNeasy Plus kit (Qiagen) according to the manufacturer’s instructions. Mice skin tissues chopped with sterile scalpel were lysed and homogenized with the QIAshredder system (Qiagen). Isolated RNA (50 ng) was subjected to qRT-PCR analysis using iTaq™ Universal SYBR^®^ Green One-Step Kit (Bio-Rad) in the 7500 Real-Time PCR System (Applied Biosystems). The comparative threshold cycle method (ΔΔCt) was used to quantify relative amounts of murine VEGF-A transcripts. Mouse GAPDH gene acts as an endogenous reference control. Primer sequences (forward and reverse, respectively) used for qRT-PCR were as follows. VEGFA: 5’-CAGGCTGCTGTAACGATGAA-3’ and 5’-TCACCGCCTTGGCTTGTCAC-3’; GAPDH: 5’-AACTTTGGCATTGTGGAAGG-3’ and 5’-ACACATTGGGGGTAGGAACA-3’.

### Statistical analysis

Statistical comparisons were performed with one-way analysis of variance (ANOVA) using GraphPad Prism 8 software. Difference between groups were considered significant when values of *P*≤0.05. To make comparisons between groups, Dunnett multiple comparison tests were performed in all applicable experiments after one-way ANOVA. Data are expressed as mean ± SEM and representative of at least two independent experiments.

## Acknowledgements

We thank Dr. Clara Curiel-Lewandrowski at the University of Arizona for generously sharing UV-exposed human skin specimens. We appreciate assistance from Mayo Clinic Pathology Research Core staff and animal facilities personnel at Mayo Clinic and The Hormel Institute, University of Minnesota.

## Funding

This study was supported by National Institutes of Health grants DK083219 (to V.M.B.), CA78383 (to D.M.), CA150190 (to D.M.), GM63904 (to S.C.E.), and CA187035 (to L.H.H.) as well as Fifth District Eagles Cancer Telethon Postdoctoral Fellowship Award (to S.K.A.), Institutional Research Grant #129819-IRG-16-189-58-IRG81 from the American Cancer Society (to L.H.H.), Austin, Minnesota “Paint the Town Pink” Award (to L.H.H.), The Mayo Foundation, and The Hormel Foundation.

## Data Availability Statement

Data pertaining to this study are included in the manuscript and supplementary information. All datasets are freely available upon request to the corresponding authors.

## Conflicts of Interest

The authors declare no conflicts of interest. Mayo Clinic has filed for patent protection for the work described in this manuscript.

## Author Contributions

S.P.H., V.M.B., S.K.A., M.O., M.S., S.H. and L.H.H. performed in vivo studies. V.M.B., S.K.A., and S.H. completed PCR experiments. S.P.H., S.K.A., A.M., and L.H.H. contributed to the completion of histology work. D.M.S. provided pathological reviews and reporting. K.P., R.A.K., P.M., D.M., and L.H.H. made significant contributions to gold nanoparticle conjugation with VEGF antibodies and the associated workflow. S.C.E. conceived and developed the original hypothesis. S.P.H., V.M.B., A.M., D.M., S.C.E., and L.H.H. performed technical troubleshooting, reviewed relevant scientific literature, critically analyzed data, and contributed to the advancement of hypotheses. S.P.H., V.M.B., S.C.E., and L.H.H. wrote the manuscript. S.P.H. and L.H.H. prepared the reported figures. S.P.H., S.K.A., A.M., S.C.E., and L.H.H. contributed to manuscript revisions. V.M.B., S.K.A., D.M., S.C.E., and L.H.H. acquired funding to complete the reported research.

---

**Supplementary Table 1:**
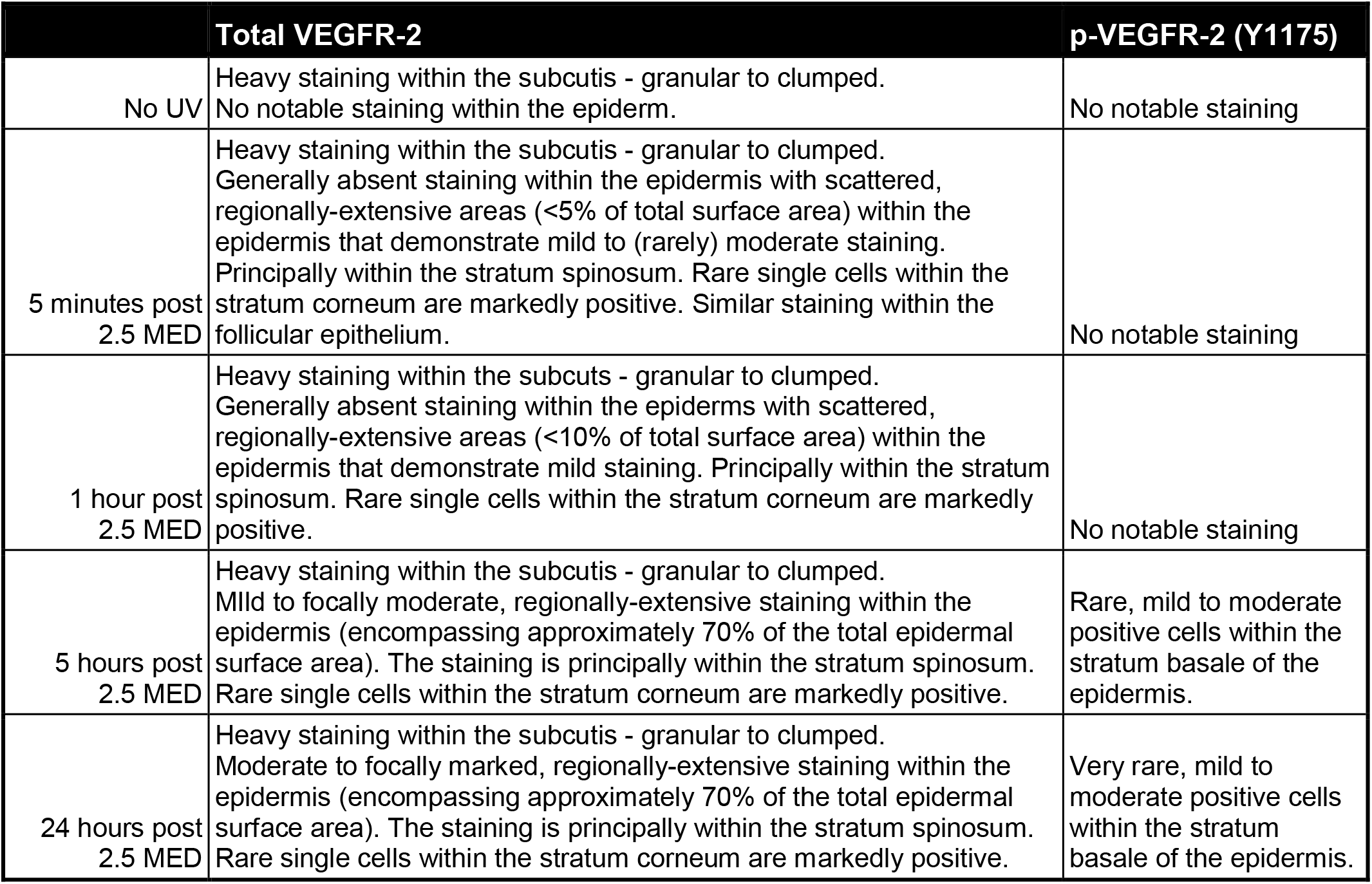
Pathological analysis of VEGFR-2 immunostaining in skin specimens derived from human subjects exposed to 2.5 MED solar simulated UV light.

**Supplementary Figure 1:**
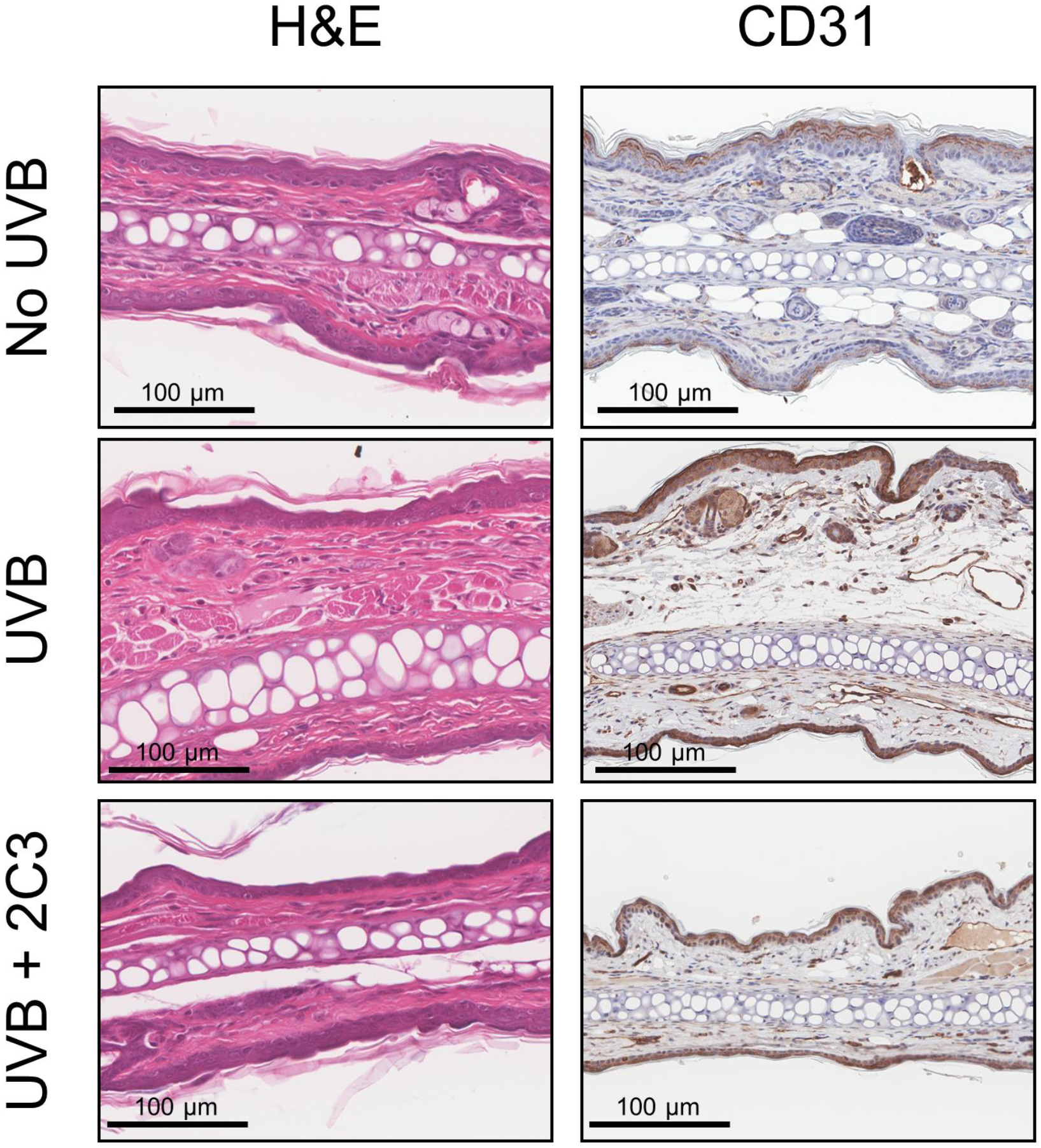
Inhibition of VEGF reduces UV-induced angiogenesis in mice. SKH1-Elite mice received a single intraperitoneal injection of 50 μg VEGF-neutralizing antibody 2C3 or negative control PBS. After 24 hours, the ears of mice were exposed to a single 1.0 MED UVB exposure of 0.144 J/cm^2^. After 48 hours, tissue was harvested and formalin-fixed, paraffin embedded specimens were immunostained for CD31. Specimens were obtained from 2 mice per group, 5 images were acquired per slide, and representative CD31 immunostaining is shown. Scale bar indicates 100 μm.

## Notes

### Competing Interest Statement

The authors have declared no competing interest.

## References

1. Guy GP, Jr., Berkowitz Z, Watson M. Estimated Cost of Sunburn-Associated Visits to US Hospital Emergency Departments. JAMA dermatology. 2017;153(1):90–2. doi: 10.1001/jamadermatol.2016.4231. PubMed PMID: 27902809; PubMed Central PMCID: PMC6057474.

2. Imokawa G. Mechanism of UVB-induced wrinkling of the skin: paracrine cytokine linkage between keratinocytes and fibroblasts leading to the stimulation of elastase. The journal of investigative dermatology Symposium proceedings. 2009;14(1):36–43. doi: 10.1038/jidsymp.2009.11. PubMed PMID: 19675551.

3. Imokawa G, Ishida K. Biological mechanisms underlying the ultraviolet radiation-induced formation of skin wrinkling and sagging I: reduced skin elasticity, highly associated with enhanced dermal elastase activity, triggers wrinkling and sagging. International journal of molecular sciences. 2015;16(4):7753–75. doi: 10.3390/ijms16047753. PubMed PMID: 25856675; PubMed Central PMCID: PMC4425048.

4. Fears TR, Scotto J, Schneiderman MA. Mathematical models of age and ultraviolet effects on the incidence of skin cancer among whites in the United States. American journal of epidemiology. 1977;105(5):420–7. PubMed PMID: 860705.

5. Watson M, Holman DM, Maguire-Eisen M. Ultraviolet Radiation Exposure and Its Impact on Skin Cancer Risk. Seminars in oncology nursing. 2016;32(3):241–54. doi: 10.1016/j.soncn.2016.05.005. PubMed PMID: 27539279; PubMed Central PMCID: PMC5036351.

6. Savoye I, Olsen CM, Whiteman DC, Bijon A, Wald L, Dartois L, Clavel-Chapelon F, Boutron-Ruault MC, Kvaskoff M. Patterns of Ultraviolet Radiation Exposure and Skin Cancer Risk: the E3N-SunExp Study. Journal of epidemiology. 2018;28(1):27–33. doi: 10.2188/jea.JE20160166. PubMed PMID: 29176271; PubMed Central PMCID: PMC5742376.

7. Detert H, Hedlund S, Anderson CD, Rodvall Y, Festin K, Whiteman DC, Falk M. Validation of sun exposure and protection index (SEPI) for estimation of sun habits. Cancer epidemiology. 2015;39(6):986–93. doi: 10.1016/j.canep.2015.10.022. PubMed PMID: 26547793.

8. Buller DB, Cokkinides V, Hall HI, Hartman AM, Saraiya M, Miller E, Paddock L, Glanz K. Prevalence of sunburn, sun protection, and indoor tanning behaviors among Americans: review from national surveys and case studies of 3 states. Journal of the American Academy of Dermatology. 2011;65(5 Suppl 1):S114–23. doi: 10.1016/j.jaad.2011.05.033. PubMed PMID: 22018060.

9. Han A, Maibach HI. Management of acute sunburn. American journal of clinical dermatology. 2004;5(1):39–47. doi: 10.2165/00128071-200405010-00006. PubMed PMID: 14979742.

10. Rastogi RP, Richa, Kumar A, Tyagi MB, Sinha RP. Molecular mechanisms of ultraviolet radiation-induced DNA damage and repair. Journal of nucleic acids. 2010;2010:592980. doi: 10.4061/2010/592980. PubMed PMID: 21209706; PubMed Central PMCID: PMC3010660.

11. Bernard JJ, Gallo RL, Krutmann J. Photoimmunology: how ultraviolet radiation affects the immune system. Nature reviews Immunology. 2019;19(11):688–701. doi: 10.1038/s41577-019-0185-9. PubMed PMID: 31213673.

12. Leung DW, Cachianes G, Kuang WJ, Goeddel DV, Ferrara N. Vascular endothelial growth factor is a secreted angiogenic mitogen. Science. 1989;246(4935):1306–9. PubMed PMID: 2479986.

13. Senger DR, Galli SJ, Dvorak AM, Perruzzi CA, Harvey VS, Dvorak HF. Tumor cells secrete a vascular permeability factor that promotes accumulation of ascites fluid. Science. 1983;219(4587):983–5. PubMed PMID: 6823562.

14. Hirakawa S, Fujii S, Kajiya K, Yano K, Detmar M. Vascular endothelial growth factor promotes sensitivity to ultraviolet B-induced cutaneous photodamage. Blood. 2005;105(6):2392–9. doi: 10.1182/blood-2004-06-2435. PubMed PMID: 15550485.

15. Yano K, Kadoya K, Kajiya K, Hong YK, Detmar M. Ultraviolet B irradiation of human skin induces an angiogenic switch that is mediated by upregulation of vascular endothelial growth factor and by downregulation of thrombospondin-1. The British journal of dermatology. 2005;152(1):115–21. doi: 10.1111/j.1365-2133.2005.06368.x. PubMed PMID: 15656811.

16. Grimes DR. Ultraviolet radiation therapy and UVR dose models. Medical physics. 2015;42(1):440–55. doi: 10.1118/1.4903963. PubMed PMID: 25563284.

17. Valejo Coelho MM, Matos TR, Apetato M. The dark side of the light: mechanisms of photocarcinogenesis. Clinics in dermatology. 2016;34(5):563–70. doi: 10.1016/j.clindermatol.2016.05.022. PubMed PMID: 27638434.

18. Lisby S, Gniadecki R, Wulf HC. UV-induced DNA damage in human keratinocytes: quantitation and correlation with long-term survival. Experimental dermatology. 2005;14(5):349–55. doi: 10.1111/j.0906-6705.2005.00282.x. PubMed PMID: 15854128.

19. Einspahr JG, Curiel-Lewandrowski C, Calvert VS, Stratton SP, Alberts DS, Warneke J, Hu C, Saboda K, Wagener EL, Dickinson S, Dong Z, Bode AM, Petricoin IE. Protein activation mapping of human sun-protected epidermis after an acute dose of erythemic solar simulated light. NPJ precision oncology. 2017;1. doi: 10.1038/s41698-017-0037-7. PubMed PMID: 29167824; PubMed Central PMCID: PMC5695572.

20. Petrova TV, Makinen T, Alitalo K. Signaling via vascular endothelial growth factor receptors. Experimental cell research. 1999;253(1):117–30. doi: 10.1006/excr.1999.4707. PubMed PMID: 10579917.

21. Simons M, Gordon E, Claesson-Welsh L. Mechanisms and regulation of endothelial VEGF receptor signalling. Nature reviews Molecular cell biology. 2016;17(10):611–25. doi: 10.1038/nrm.2016.87. PubMed PMID: 27461391.

22. Abhinand CS, Raju R, Soumya SJ, Arya PS, Sudhakaran PR. VEGF-A/VEGFR2 signaling network in endothelial cells relevant to angiogenesis. Journal of cell communication and signaling. 2016;10(4):347–54. doi: 10.1007/s12079-016-0352-8. PubMed PMID: 27619687; PubMed Central PMCID: PMC5143324.

23. Man XY, Yang XH, Cai SQ, Yao YG, Zheng M. Immunolocalization and expression of vascular endothelial growth factor receptors (VEGFRs) and neuropilins (NRPs) on keratinocytes in human epidermis. Molecular medicine. 2006;12(7-8):127–36. doi: 10.2119/2006-00024.Man. PubMed PMID: 17088944; PubMed Central PMCID: PMC1626599.

24. Zhu JW, Wu XJ, Luo D, Lu ZF, Cai SQ, Zheng M. Activation of VEGFR-2 signaling in response to moderate dose of ultraviolet B promotes survival of normal human keratinocytes. The international journal of biochemistry & cell biology. 2012;44(1):246–56. doi: 10.1016/j.biocel.2011.10.022. PubMed PMID: 22062947.

25. Brauchle M, Funk JO, Kind P, Werner S. Ultraviolet B and H2O2 are potent inducers of vascular endothelial growth factor expression in cultured keratinocytes. The Journal of biological chemistry. 1996;271(36):21793–7. doi: 10.1074/jbc.271.36.21793. PubMed PMID: 8702976.

26. Krutmann J, Grewe M. Involvement of cytokines, DNA damage, and reactive oxygen intermediates in ultraviolet radiation-induced modulation of intercellular adhesion molecule-1 expression. The Journal of investigative dermatology. 1995;105(1 Suppl):67S–70S. PubMed PMID: 7616000.

27. Trompezinski S, Pernet I, Schmitt D, Viac J. UV radiation and prostaglandin E2 up-regulate vascular endothelial growth factor (VEGF) in cultured human fibroblasts. Inflammation research : official journal of the European Histamine Research Society [et al]. 2001;50(8):422–7. doi: 10.1007/PL00000265. PubMed PMID: 11556523.

28. Mukherjee P, Bhattacharya R, Wang P, Wang L, Basu S, Nagy JA, Atala A, Mukhopadhyay D, Soker S. Antiangiogenic properties of gold nanoparticles. Clinical cancer research : an official journal of the American Association for Cancer Research. 2005;11(9):3530–4. doi: 10.1158/1078-0432.CCR-04-2482. PubMed PMID: 15867256.

29. Yano K, Kajiya K, Ishiwata M, Hong YK, Miyakawa T, Detmar M. Ultraviolet B-induced skin angiogenesis is associated with a switch in the balance of vascular endothelial growth factor and thrombospondin-1 expression. The Journal of investigative dermatology. 2004;122(1):201–8. doi: 10.1046/j.0022-202X.2003.22101.x. PubMed PMID: 14962109.

30. Rawlings JS, Rosler KM, Harrison DA. The JAK/STAT signaling pathway. Journal of cell science. 2004;117(Pt 8):1281–3. doi: 10.1242/jcs.00963. PubMed PMID: 15020666.

31. Hao D, Wen X, Liu L, Wang L, Zhou X, Li Y, Zeng X, He G, Jiang X. Sanshool improves UVB-induced skin photodamage by targeting JAK2/STAT3-dependent autophagy. Cell death & disease. 2019;10(1):19. doi: 10.1038/s41419-018-1261-y. PubMed PMID: 30622245; PubMed Central PMCID: PMC6325150.

32. Alam SK, Astone M, Liu P, Hall SR, Coyle AM, Dankert EN, Hoffman DK, Zhang W, Kuang R, Roden AC, Mansfield AS, Hoeppner LH. DARPP-32 and t-DARPP promote non-small cell lung cancer growth through regulation of IKKalpha-dependent cell migration. Commun Biol. 2018;1:43. doi: 10.1038/s42003-018-0050-6. PubMed PMID: 29782621; PubMed Central PMCID: PMC5959014.

